# Uncovering *Staphylococcus aureus* genes with roles in pathogenicity by silkworm infection model

**DOI:** 10.1101/714725

**Authors:** Atmika Paudel, Hiroshi Hamamoto, Suresh Panthee, Yasuhiko Matsumoto, Kazuhisa Sekimizu

## Abstract

The regulatory network of virulence factors production by *Staphylococcus aureus*, an opportunistic pathogen, is incompletely understood, and the functions of many uncharacterized genes in its genome remain to be uncovered. We screened 380 function unassigned genes disrupted mutants of the community-acquired methicillin-resistant *S. aureus* USA300 for pathogenicity using silkworms and identified 11 strains with reduced silkworm killing ability. Nine out of the 11 strains displayed reduced virulence in the mouse model as evidenced by reduced colony-forming units in organs of the infected mice. Three of the identified gene-disrupted mutants had reduced hemolytic activity, one among the three also had reduced proteolytic activity and pigment production. These results suggest that silkworm model could identify the genes required for virulence in the mouse model. The newly identified genes involved in virulence in this study facilitates the further understanding of the pathogenicity of *S. aureus*.

**Importance:** We performed a large scale screening of mutants of *Staphylococcus aureus* with disruption in function unassigned genes using silkworm infection model and identified eleven genes required for full virulence in silkworm. Nine of the eleven genes were involved in virulence in mice and were previously not known to aggravate virulence of *S. aureus*. The results suggest that silkworm model is suitable for quantitative measurement of virulence, which is shared between silkworms and mammals.

## Introduction

*Staphylococcus aureus* is a commensal and an opportunistic pathogen that is responsible for ailments ranging from skin infection to deep tissue infection, endocarditis, and bacteremia(1). With an increasing rise in the resistance of methicillin-resistant *S. aureus* (MRSA), the appearance of vancomycin-intermediate *S. aureus* (VISA)(2, 3), and vancomycin-resistant *S. aureus* (VRSA)(4–7), *S. aureus* has become one of the leading causes of morbidity and mortality and has been categorized as a threat among antimicrobial-resistant strains by the World Health Organization(8) and the Centers for Disease Control and Prevention(9). The ability of *S. aureus* to rapidly infect host is due to the expression and release of many virulence factors such as cytolysins, hemolysins, leukocidins, coagulases, adhesins, proteases, nucleases, enterotoxins, lipases, exfoliative toxins, and immune-modulatory factors, and cell surface associated proteins including Protein A and fibrinogen-, fibronectin- and collagen-binding proteins(10–15). Although some of the pathways of virulence factor expression and regulation such as *sarA*, *agr*, *srrAB*, *saeRS*, *ArlRS*,(16) in *S. aureus* as well as the factors acting via one or more of these pathways(17–19) are studied, the understanding of the production and regulation of virulence factors in *S. aureus* is through a multifaceted network that is still obscure and many genes with unknown functions in *S. aureus* genome with roles in virulence and pathogenicity remain to be uncovered.

Since virulence is the degree of pathogenicity of a microorganism that is mostly affected by host environment and host-pathogen interaction, a suitable model that addresses host-pathogen interactions and mimics the clinical condition of infection is required for the understanding of the pathogenicity of *S. aureus* on a molecular level. Use of mouse models for screening of virulence factors is impractical due to high costs and the associated ethical issues. Therefore, utilization of a model that has less ethical issues, resembles a mouse systemic infection model, and allows quantitative analysis of virulence is desired. Here, we used silkworms (*Bombyx mori*) to uncover the roles of uncharacterized genes of *S. aureus* in pathogenicity. We have previously shown that injection of *S. aureus* kills the silkworms(20) and activates innate immunity(21). We can evaluate the therapeutic effectiveness of antibiotics using silkworm infection model by comparison of effective dose fifty (ED_50_) values of clinically used antibiotics between silkworms and mammals(22). On the basis of these advantages, we used silkworms to discover various antibacterial agents against *S. aureus* infections such as lysocin E from *Lysobacter sp.* RH2180-5 (23–25), GPI0363(26), compound 5(27) and iminothiadiazolo-pyrimidinone derivatives(28). Additionally, silkworms have large enough body size to allow administration of precise inoculum and dosage, are economical, easy to handle, and associated with less ethical issues as well as less biohazard potential(29, 30). The number of bacterial cells required to kill silkworms can be quantitatively determined by the administration of precise inoculum of the bacteria. This allows a direct and easy comparison of the degree of pathogenicity among bacterial strains from their respective lethal dose fifty (LD_50_) values. We have previously identified novel *S. aureus* virulence factors using silkworms such as CvfA, CvfB, CvfC(31), SarZ(32) and SA1684(33) that have roles in pathogenicity in mice models. In this regard, the use of silkworm as a model is the unique way for the identification of novel virulence factors of *S. aureus* that can have roles in pathogenicity to mammals. Since community-acquired MRSA (CA-MRSA) USA300 is hypervirulent compared to the majority of the hospital-acquired MRSAs(34), we used USA300 strain for screening in this study. Here, we performed the first large-scale and quantitative screening of the gene-disrupted mutants of USA300 JE2 from the Nebraska Transposon Mutant Library (NTML)(35) using silkworms. We identified 11 mutants with reduced virulence in silkworms, among which nine had reduced virulence in mice. To our knowledge, this is the first report that uncovers the roles of these genes in aggravating the infection.

## Results

### Identification of novel S. aureus genes with roles in pathogenicity using silkworms

Previous attempts of evaluation of virulence genes in *S. aureus* by using silkworms have been performed in small scales and with methicillin-susceptible *S. aureus* (MSSA) strains such as RN4220(31) and NCTC8325-4(36). In this study, we performed a large-scale screening of a mutant library of a CA-MRSA USA300 LAC JE2 strain to find genes required for full virulence in silkworms for the first time. At first, a total of 380 gene-disrupted mutants whose gene products were hypothetical proteins and were not tested previously using silkworms(31) were selected for the study. As silkworms can be injected with accurate volumes of inoculum by using syringes, the primary screening was performed by injection of same dilutions (ten-fold) of the overnight culture of the strains and observing the survival of the infected silkworms. We obtained 28 strains in the primary screening that did not kill silkworms when the wild-type strain killed all the silkworms (Figure 1a, **Supplementary Table S1**). Next, we compared the lethal dose fifty (LD_50_) values, the bacterial dose that kills half of the silkworms, after injecting serial dilutions of the cultures into the silkworm hemolymph. Counting the colony-forming units (CFU) of the injected bacteria was performed separately. Out of 28, 12 strains showed ≥ 2 fold higher LD_50_ values than that of the wild-type (**Supplementary Table S1**). We, then, selected these 12 strains for further examination and found that 11 showed higher LD_50_ values with statistical significance in silkworms compared to that of the wild-type (Figure 1a,b). The feasibility of accurate inoculum administration and quantitative analysis can be attributed to the discovery of previously unidentified genes with roles in pathogenicity.

**Figure 1:**
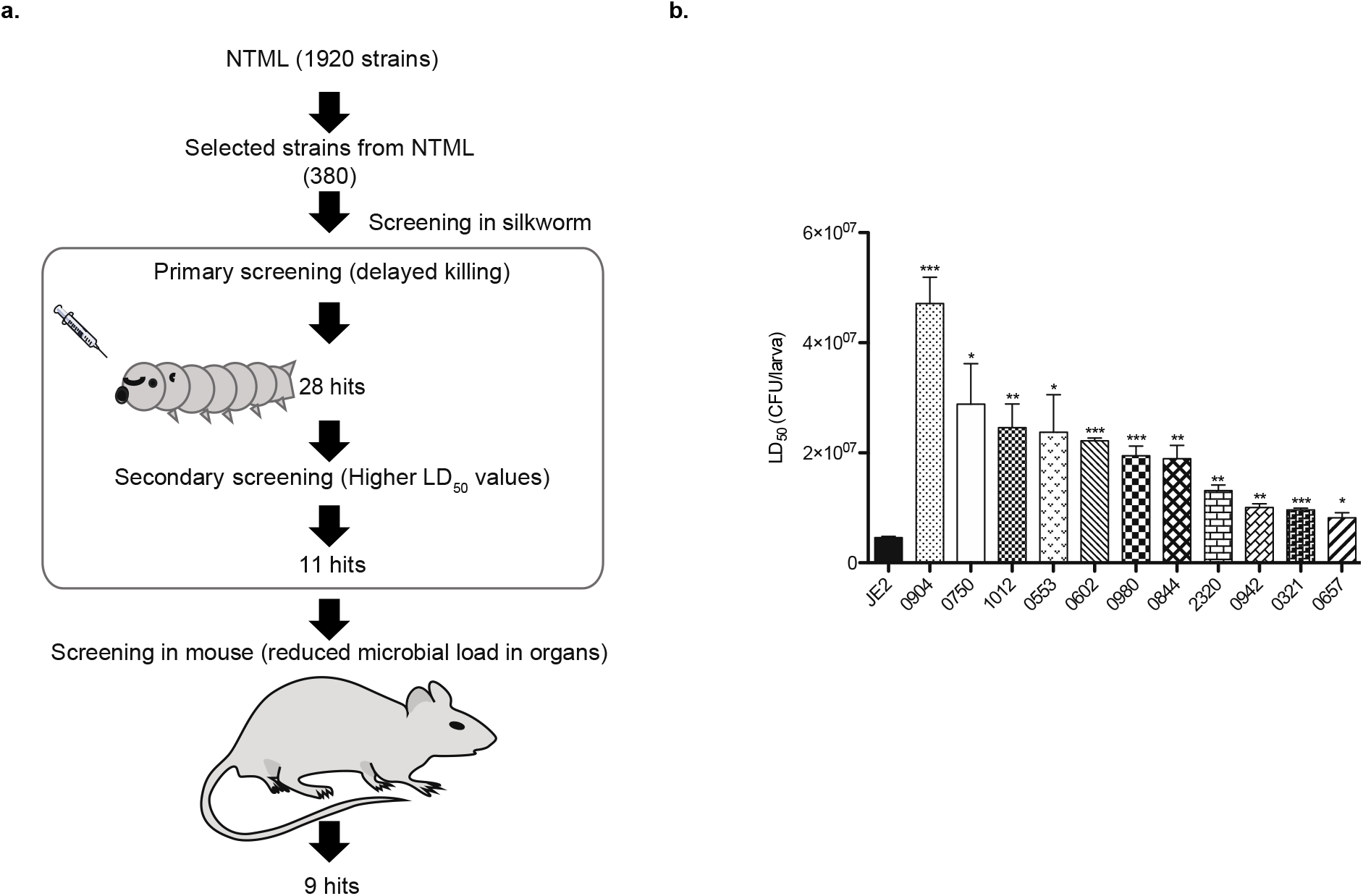
Screening and identification of novel candidate virulence factors of *S. aureus* using silkworms. a: Screening strategy and results of screening. b: Strains exhibiting reduced pathogenicity to silkworms compared to the wild-type. Overnight cultures of *S. aureus* gene-disruptant mutants were serially diluted and injected into the silkworm hemolymph (n=5), and survival was determined 30h post infection. LD_50_ values were determined using logistic regression analysis using R-studio Program. Data are means ± S.E.M of three independent experiments, analyzed by Student’s unpaired *t*-test and significant differences compared with the wild-type are indicated by asterisks (\**p* ≤ 0.05, ** *p* ≤ 0.01, \*\*\**p* ≤ 0.001).

### Pathogenicity of the identified mutants in the mouse infection model

We evaluated the pathogenicity of the 11 strains in the mouse systemic infection model by examining the microbial burden in organs-heart and kidney of the infected mice. Nine out of 11 strains had significantly reduced microbial survival in at least one of the organs (Figure 2a, b) indicating that more than 80% of the virulence factors identified using silkworms exert virulence in mammals. This finding shows that the silkworm infection model efficiently identifies bacterial factors with roles in pathogenicity to mammals.

**Figure 2:**
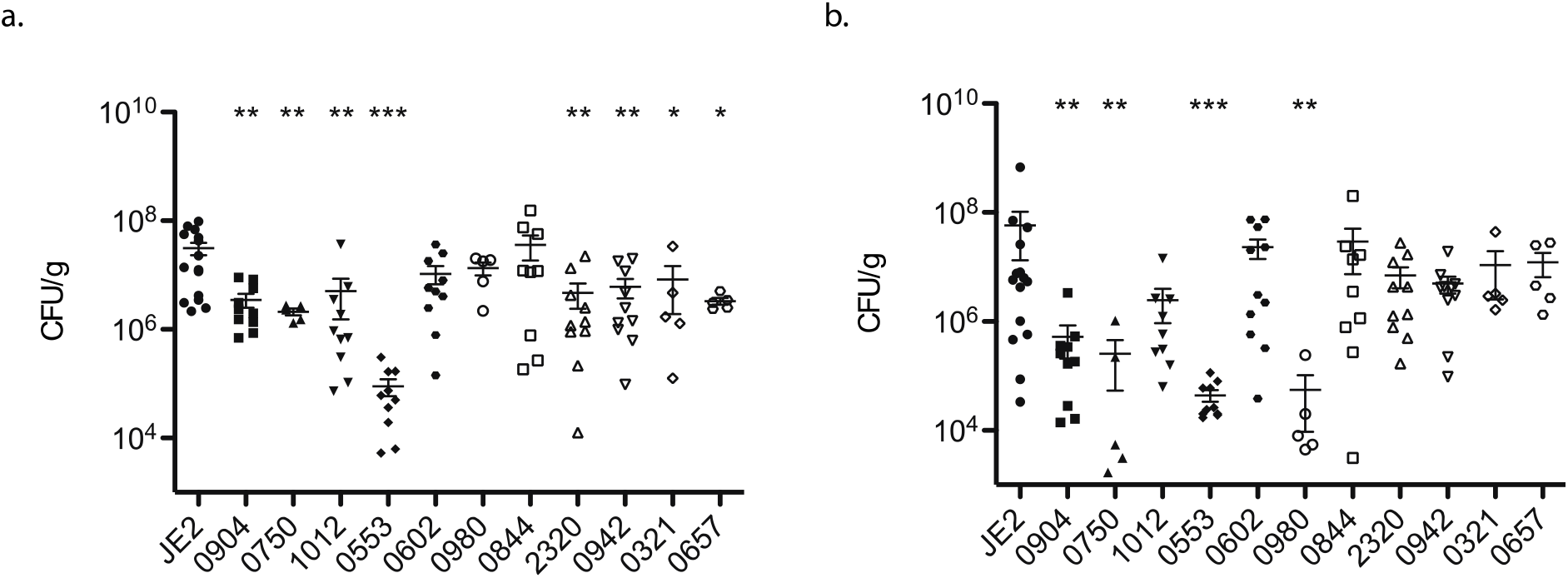
Microbial burden in kidney (a) and heart (b). Mice were injected intravenously with the strains into the tail vein and bacteria recovered from the isolated organs (kidney and heart) was counted. Each symbol represents data obtained from one animal. Data are shown as mean ± S.E.M, analyzed by the Mann–Whitney U-test and significant differences compared to the wild-type are indicated by asterisks (\**p* ≤ 0.05, ** *p* ≤ 0.01, and \*\*\**p* ≤ 0.001). Average injected CFU: JE2: 1.2×10^8^; 0553: 1.2×10^8^; 1012: 9.9×10^7^; 0750: 1.3×10^8^; 0904: 1.2×10^8^; 0657: 1.3×10^8^; 2320: 1.3×10^8^, 0942: 1.2×10^8^; 0321: 1.3×10^8^; 0602: 1×10^8^; 0980: 1.4×10^8^; 0844: 2×10^8^.

While comparing the microbial burden of the above mentioned nine strains with that of the wild-type strain, three strains (*0553::Tn*, *0750::Tn* and *0904::Tn*) had a lower microbial burden in both the organs; five strains (*1012::Tn*, *0657::Tn*, *2320::Tn*, *0942::Tn* and *0321::*Tn) had a reduced survival in kidney while their survival in heart was indistinguishable; and one strain (*0980::Tn*) displayed a lower survival in heart and an indistinguishable difference in kidney. The findings indicate that the patterns of colonization efficiencies in each mice organ were different between the gene-disrupted strains.

### In vitro phenotypes of the mutants

We examined the growth rates of all the mutants grown *in vitro* in TSB medium at 37°C. We found that the doubling times of the mutants were not significantly different from that of the wild-type (**Supplementary Figure S1**) suggesting that the differences in pathogenicity could not be attributed to differences in growth rates of the strains. To further check whether the phenomena of low pathogenicity is related to the reduction in toxin production, we compared the hemolytic and proteolytic activities of the mutants with those of the wild-type. We found that three strains *0904::Tn*, *0750::Tn* and *0657::Tn* had reduced hemolytic activity as determined by a clear zone surrounding the colony in sheep-blood (Figure 3) and that *0904::Tn* had reduced proteolytic activity as evaluated by a clear zone surrounding the colony in skim milk agar plates (Figure 3). Besides, the *0904::Tn* strain had a reduced pigment production capacity on the sheep blood agar plate (Figure 3). The reduced pigment and protease production of *0904::Tn* were in accordance to previous reports(35, 37).

**Figure 3:**
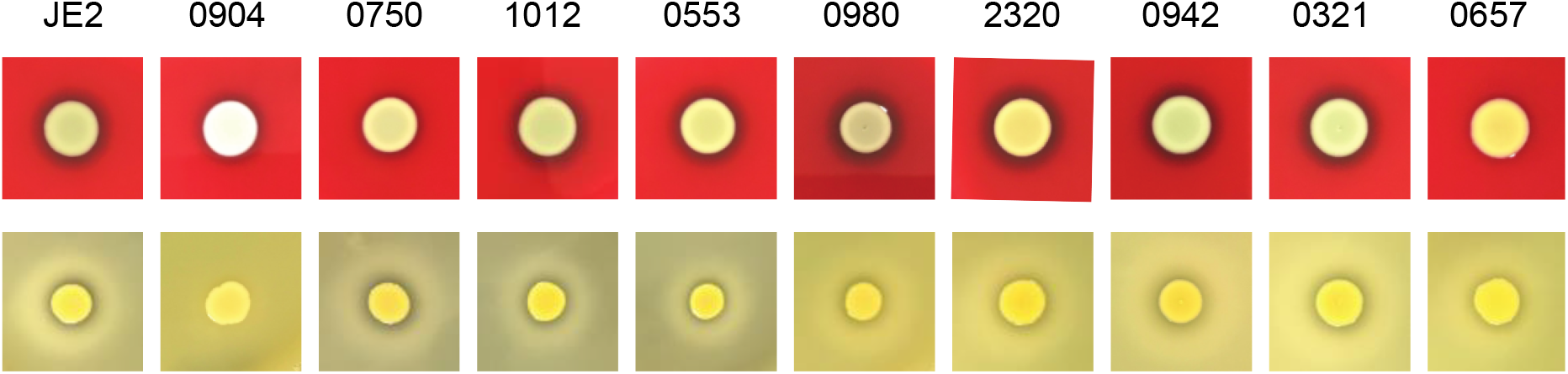
Hemolytic and proteolytic activities of the strains. Two microliters of the overnight cultures of the strains were spotted on sheep blood agar (upper panel) and TSB+ 3.3% skim milk agar plates (lower panel) and incubated at 37°C overnight. Data are representative of three independent experiments.

Among the analyzed strains, the LD_50_ value in silkworm was the highest for the *0904::Tn* strain which could not lyse sheep red blood cells and casein, and had a reduced pigment production capacity suggesting that the loss of pathogenicity of this strain could be explained at least in part by the reduced toxin production capacity. For the strains with the indistinguishable change in the toxin production capacity, we assume that these genes regulate virulence in *S. aureus* via pathways other than toxin production and that the loss of pathogenicity could be a result of various host-pathogen interplay. This suggested that the candidates identified in this study would have been missed in the *in vitro* screens, highlighting the importance of *in vivo* screening system for the screening of genes involved in virulence.

To understand the properties of these novel virulence factors, we performed a bioinformatic analysis and predicted the functional domains of these proteins. First, based on the description in the GenBank, we found that all but two were categorized as hypothetical proteins. Next, we obtained the complete genome sequences of 328 *S. aureus* strains from GenBank and analyzed the orthologous proteins by Orthofinder 2.2.6(38) as explained previously(39). We found that the nine proteins identified in this study were highly conserved (detected in nearly 97% strains) among different strains of *S. aureus* while eight strains did not harbor the homologs for five of the identified virulence factors: SAUSA300_0553, SAUSA300_0904, SAUSA300_0942, SAUSA300_1012, and SAUSA300_2320 (**Supplementary Table S2**). Furthermore, we found that four out of the nine proteins were predicted to be located in the cytoplasm and three in the cytoplasmic membrane; four were predicted to consist of transmembrane helicases; one was predicted to be a secretory protein; and six were categorized to be proteins with either no known homologous superfamily or having an uncharacterized domain (Table 1). Detailed analysis of the molecular mechanism of how these proteins act as virulence factors will be the next step in our research.

**Table 1:** The functional domains of the proteins with roles in pathogenicity. The analyses were performed using: PSORTB Ver 3.0.2(44); TMPred (minimum length: 14, maximum: 33)(45); LnSignal(46); and Interpro(47) for the prediction of location, transmembrane helicase, secretory protein and homologous families/superfamilies, respectively.

| SAUSA300_ | AA length | Definition (GenBank) | Location | Transmembrane helix | Secretory protein | Homologous family (Interpro ID) |
| --- | --- | --- | --- | --- | --- | --- |
| 0904 | 121 | Protozoan/cyanobacterial globin family protein | Unknown | Yes, N-terminus outside | Non-secretory | Truncated hemoglobin (IPR001486) |
| 0750 | 314 | Hypothetical protein | Cytoplasm | No | Non-secretory | Sporulation regulator WhiA-like (IPR033981) |
| 1012 | 91 | Hypothetical protein | Cytoplasm | No | Non-secretory | Uncharacterised protein family UPF0358 (IPR009983) |
| 0553 | 120 | Hypothetical protein | Unknown | No | Non-secretory | Protein of unknown function DUF1806 (IPR014934) |
| 0980 | 431 | Membrane protein | Cytoplasmic membrane | Yes, N-terminus inside | Secretory | None |
| 2320 | 143 | Hypothetical protein | Cytoplasmic membrane | Yes, N-terminus outside | Non-secretory | Protein of unknown function DUF2871 (IPR021299) |
| 0942 | 96 | Hypothetical protein | Cytoplasmic membrane | Yes, N-terminus inside | Non-secretory | None |
| 0321 | 275 | Hypothetical protein | Cytoplasm | No | Non-secretory | Alpha/Beta hydrolase fold (IPR029058) |
| 0657 | 214 | Hypothetical protein | Cytoplasm | No | Non-secretory | None |

## Discussion

The success of *S. aureus* to infect host depends on its ability to evade the immune system and acquire nutrition from the host. The genomic analysis of multiple *S. aureus* indicates the presence of several genes encoding for hypothetical or uncharacterized proteins. Some of these genes are conserved among many *S. aureus* strains indicating that they might serve as a tool to compete in the host environment. In our study, we found that some of these conserved uncharacterized proteins were, in fact, involved in *S. aureus* pathogenicity as virulence factors.

Several virulence factors and their regulators have been exploited as targets for screening and identification of inhibitors; however, the intricate multidimensional nature of virulence factors expression and regulation makes it difficult to select a single virulence factor as a target. Moreover, the expression of virulence factors is largely dependent upon the host environment(40) and stage of infection. Therefore, for target-based screening, it is important to select the virulence factor(s) that are expressed while infecting a host and be responsible for full virulence. We used silkworms for this purpose. Since silkworms are easy to rear in a small space, easy to handle, and special injection techniques are not required, it is possible to inject hundreds of silkworms within an hour. Most importantly, less ethical concerns associated with its use and feasibility of quantitative administration of precise inoculum/dose make the silkworm an excellent model host for screening purposes. The established infection models, activation of innate-immunity by pathogen invasion, treatment by clinically used antimicrobial agents, and presence of basic pharmacokinetic pathways common to mammals advocate that silkworms are suitable for the study of host-pathogen interactions.

Here, we employed a simple model host-based approach using silkworms and uncovered nine novel genes that are required for full virulence of *S. aureus*. The success relied on the quantitative screening using silkworms resulting in a high degree of correlation between the virulence in silkworm and mice. Microbes such as *S. aureus* also take advantage of the host’s acquired immunity to establish infection(41), and silkworms only have an innate immune system. Therefore, the use of silkworm model can identify virulence factors that trigger innate immunity and not the acquired immunity.

The disruption of the identified genes had different effects on the tissue-specific colonization of *S. aureus* in mice likely due to the ability of *S. aureus* to generate physiologically distinct phenotypes for survival on a particular host environment(42) as well as the ability to activate and respond to the different immune cells recruited to specific organ by the host in response to the infection(43).

The identified genes have not been previously identified to have roles in aggravating virulence. However, in contrast to our result, a recent study(37) showed that retro-orbital injection of *0904::Tn* resulted in increased colonization in the C57BL/6J mouse kidney. The differences might have been observed owing to the differences in the infection models and the genetic backgrounds of the mice used. Our survival assay in silkworm systemic infection model clearly indicates that *0904::Tn* has reduced virulence, which is consistent with the reduced colonization of the mutant in the organs of a mouse systemic infection model.

In summary, by the identification of novel virulence factors, our study facilitates the understanding of host-pathogen interaction and the complex network of virulence factors production and regulation of *S. aureus*. The gene products identified in this study could serve as the targets for the treatment of infections by *S. aureus*. This study further justifies the use of silkworm model for the discovery of therapeutically effective antibiotics and screening novel virulence factors using silkworm model, which can be utilized to other pathogenic microorganisms besides *S. aureus*.

## Materials and Methods

### Mutant library and bacterial growth conditions

The transposon mutant library of *S. aureus* USA300 strain JE2 referred to as-Nebraska Transposon Mutant Library Screening Array was kindly provided by BEI resources (NR-48501). The library consists of gene disruptants of 1920 genes. Single colonies of the wild-type and the gene-disrupted mutants were isolated in tryptic-soy broth (TSB; Becton Dickinson and Company, Franklin Lakes NJ, USA) agar plates with supplementation of 5 µg/ml erythromycin (Wako Pure Chemicals, Tokyo, Japan) for the mutants. A single colony was then inoculated in 5 ml of TSB or TSB supplemented with erythromycin and grown overnight by shaking at 37°C to prepare the full growth.

### Silkworm rearing and infection assay

Silkworms were raised as previously described(26). The fifth instar larvae were fed an antibiotic-free artificial diet for one day, and 50 µl of each dilution of the strains was injected into the hemolymph of larvae (n=10) using a 1-ml syringe (Terumo Corporation, Tokyo, Japan) equipped with a 27-gauge needle (Terumo Corporation). The silkworms were then kept at 27°C, and survival was observed.

### Screening of gene-disrupted mutants for reduced pathogenicity in silkworms

The screening was performed in two stages:

#### Primary screening

The full growth of the wild-type JE2 and the selected 380 gene-disrupted mutants was diluted tenfold with sterile 0.9% NaCl (Otsuka Normal Saline, Otsuka Pharmaceutical Factory, Inc., Tokyo, Japan) and injected into silkworms. Strain with mutation in *cvfB* gene, SAUSA300_1284 was used as a positive control(31). Those strains with reduced silkworm killing ability under the following criteria were selected as hits: survival of at least 50% silkworms when all silkworms injected with the wild-type died and all silkworms injected with the vehicle survived.

#### Secondary screening

The overnight culture of the strains was subjected to two-fold serial dilution with sterile 0.9% NaCl (Otsuka Normal Saline, Otsuka Pharmaceutical Factory) and injected into the hemolymph of silkworm larvae (n=5). Survival of the larvae was observed. The colony-forming units (CFU) of the injected strains was counted by diluting the bacterial suspension, spreading onto Mueller-Hinton Broth (MHB; Becton Dickinson and Company) agar plates and incubating for overnight at 37°C. The lethal dose fifty (LD_50_) values of each strain were determined from the CFU and survival of silkworms. Strains that showed significantly higher LD_50_ values than that of the wild-type were selected for further experiments.

### Pathogenicity of the strains in mice

All mouse protocols were approved by the Teikyo University Animal Ethics Committee. Overnight cultures of the strains were diluted 100 fold and incubated at 37°C with shaking for 16 hours. The resulting cultures were then centrifuged, washed with Phosphate Buffered Saline (PBS; Wako Pure Chemicals), and suspended in PBS to adjust the turbidity (A_600_) to 0.7 using a spectrophotometer (UV-1280 Shimadzu Corp., Kyoto, Japan). Mice (ICR, female, eight weeks, Charles River Laboratories Japan Inc., Kanagawa, Japan) were injected with 200 µl of the prepared suspension of strains intravenously into the tail vein. The injected suspensions were diluted, spread on agar plates, and appeared colonies were counted to obtain the injected CFU of each strain. After 24 hours, organs (kidney and heart) of the infected mice were isolated, suspended in PBS and homogenized, diluted, and spread on agar plates. Appeared colonies were counted after overnight incubation of the plates at 37°C.

### Determination of hemolytic and proteolytic activities

The strains were cultured in TSB or TSB supplemented with erythromycin at 37°C 200 rpm overnight. Two microliters of the overnight cultures were spotted on sheep-blood agar plates (Nissui Pharmaceutical Co. Ltd., Tokyo, Japan), and TSB agar plates supplemented with 3.3% skim milk to determine hemolytic and proteolytic activity, respectively. The plates were then dried under the clean bench and incubated at 37°C overnight. The activity was determined by the appearance of the clear zone surrounding the grown bacteria.

## Acknowledgments

This work was supported by JSPS KAKENHI Grant Number JP15H05783 (Grant-in-Aid for scientific research (S)) to KS; and in part by a JSPS KAKENHI Grant Number JP19K16653 (Grant-in-Aid for Early-Career Scientists) to AP, JSPS KAKENHI Grant Number JP19K07140 (Grant-in-Aid for scientific research) to HH, and JSPS KAKENHI Grant Number JP17F17421 (Grant-in-Aid for JSPS Fellows) to SP and KS. The authors would like to acknowledge Genome Pharmaceuticals Institute Co., Ltd. for the help in the primary screening for pathogenicity in silkworms.

